# Optimal combined chemotherapeutic and antibiotic treatment for cancer in the presence of saboteur bacteria

**DOI:** 10.64898/2026.11.13.766354

**Authors:** Anna Geretovszky, Gergely Röst

## Abstract

We consider a mathematical model of cancer dynamics under chemotherapeutic treatment in the presence of bacteria capable of metabolizing the chemotherapeutic agent, thereby compromising therapeutic efficacy. To counteract this effect, we investigate the introduction of antibiotic co-therapy aimed at eradicating the bacterial population or at least mitigating its negative impact on treatment outcomes. The underlying dynamics are formulated as a system of four non-linear ordinary differential equations, with the time-varying dosages of both drugs treated as control variables. Applying optimal control theory, we explore which combined treatment strategy leads to the best outcome, and how it is influenced by the model parameters. Convergence to the optimal control is achieved by employing a relaxation method within the forward-backward sweep algorithm. Furthermore, we analyze a discrete-dose variant of the optimal control problem, providing a more clinically realistic framework for scheduled drug administration.

## 1 Introduction

Although significant progress has been made over the years, curing cancer is still one of the biggest medical challenges. Among other factors, resistance occurring during chemotherapy treatments plays a major role, since the previously effective drugs can no longer work adequately. There can be a number of reasons for this, some of which are still unknown, but the wide variety of causes includes natural selection among tumour cells, cellular plasticity, DNA reparability and even microvesicles ([1, 2, 3, 4, 5]). Research in the past few years has shown that microorganisms can also influence the effectiveness of chemotherapy treatments. The article [6] describes how certain species of bacteria can metabolise chemotherapy drugs and prevent them from having an anti-cancer effect, i.e. the tumour becomes resistant with the help of the bacteria. In such cases, it is advisable to supplement the chemotherapy treatment with a course of antibiotics to counteract the sabotaging effect of the bacteria.

Earlier, in [7], we constructed a mathematical model to represent this biological mechanism. To allow us to perform a global analysis of the dynamics of this system, we assumed that both the chemotherapeutic and antibiotic drugs are administered continuously in constant doses during the course of the treatment. Although it can be realistic under certain conditions (during in vitro experiments or IV therapies), it is a natural idea to investigate the possibility of time-dependent (but still continuous) drug doses. Our aim is to find an optimal combined treatment of the two drugs, which minimizes the harmful effects, but also eradicates the tumour. For this, we use Potryagin’s minimum principle and a relaxed Forward-Backward Sweep Method. Based on simulations, we inspect how the variation of certain parameters affect the optimal solutions. We also consider a discretized version of the continuous optimal drug doses – which resembles to taking pills – and see where it can lead us.

## 2. Differential equation model

### 2.1 Construction

The model describing the phenomenon is the non-linear differential equation system

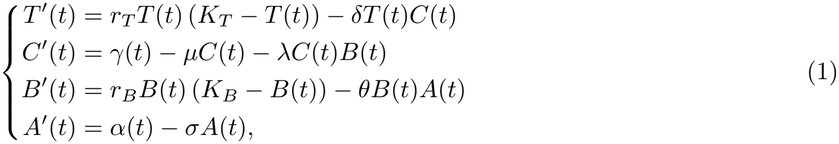

where *T* and *B* denote the sizes of the tumour and the bacterial population, respectively, and *C* and *A* denote the concentrations of the chemotherapeutic and the antibiotic drugs, respectively, as functions of time. We have *γ* as the dose of the chemotherapy, and *α* as the dose of the antibiotics, also as functions of time. A detailed breakdown of how and why this equation system was constructed can be found in [7]. The nonnegative parameters used to set up the system of equations are summarized in Table 1. In this paper, our goal is to interpret *γ* and *α* as controls in this system, and find the optimal control that gets rid of the tumour the most effective and least harmful way.

**Table 1:** Variables and parameters in the equation system (1).

| variable/param. | meaning of the variable/parameter |
|---|---|
| *T* | size of the tumour |
| *C* | concentration of the chemotherapeutic drug |
| *B* | size of the bacterial population |
| *A* | concentration of the antibiotics |
| γ | dose of the chemotherapeutic drug |
| α | dose of the antibiotics |
| *r*<sub>*T*</sub> | growth rate of the tumour |
| *K*<sub>*T*</sub> | carrying capacity wrt. the tumour |
| δ | rate of death by the chemotherapeutic drug |
| μ | absorption or elimination of the chemotherapeutic drug |
| λ | rate of chemotherapeutic drug metabolized by the bacteria |
| *r*<sub>*B*</sub> | growth rate of the bacterial population |
| *K*<sub>*B*</sub> | carrying capacity wrt. the bacteria |
| θ | rate of death of bacteria by the antibiotics |
| σ | absorption or elimination of the antibiotics |

### 2.2 Analysis

Using constant *γ*(*t*) *≡γ* and *α*(*t*) *≡α* functions, we analyzed the system (1) in [7]. It is well-defined, so for nonnegative initial values there always exists a unique bounded nonnegative solution. We constructed three combined threshold parameters, which allowed us to identify the possible outcomes of the treatment. The system has four different types of equilibria, characterized by the extinction or persistence of the tumour and the bacterial population. We showed global asymptotic stability for all four scenarios, and determined what kind of treatment is needed for the optimal outcome. Simulations were run to confirm the four possible outcomes, as well as the situations in which they are achieved. Between the different cases, we examined the types of bifurcations. We also fitted a specified version of the model to real data, showing that it indeed accurately describes the processes and interactions that take place.

In this work, we consider piecewise continuous, time-dependent functions for *γ*(*t*) and *α*(*t*), and therefore it is important to establish that the system (1) remains well-defined for such controls.

#### Proposition 2.1

(Well-posedness). *For nonnegative initial values and piecewise continuous functions γ*(*t*) *≤ γ*_*max*_, *α*(*t*) *≤ α*_*max*_, *where γ*_*max*_, *α*_*max*_ *∈* ℝ, *the equation system* (1) *has unique, nonnegative solutions, that are bounded from above. The solutions continuously depend not only on the initial values, but also on γ*(*t*) *and α*(*t*).

**Proof** . The first part follows similarly to the argument presented in Proposition 2.1 in [7], with the additional assumption that *γ*(*t*) and *α*(*t*) are bounded from above (their nonnegativity is naturally imposed by their biological interpretation). However, we must make a slight modification because the Picard-Lindelof theorem cannot be applied directly due to possible points of discontinuity in the controls. But on the time intervals between consecutive discontinuity points, the right-hand side of the system is continuous in *t*, so the Picard-Lindelöf theorem is applicable, and unique solutions exist on each such interval. Then we can concatenate these partial solutions and obtain that the equation system (1) indeed has a unique, piecewise continuously differentiable solution on the whole time interval.

Again, for every subinterval where *γ*(*t*) and *α*(*t*) are continuous, *f* is continuous and locally Lipschitz continuous in the second argument, so the solutions depend continuously on the initial conditions. Moreover, they also depend continuously on the functions *γ*(*t*) and *α*(*t*) (see Theorem 2.8 in [8]).

It is worth noting that, in the investigations presented later in Section 4.1, we will also consider cases in which no explicit finite upper bounds are imposed on these controls (in particular, infinity is used as the upper bound in the corresponding numerical simulations); we may therefore allow arbitrarily large control values theoretically. However, should an optimal control take an unbounded or unrealistically large value, such a solution would be discarded on practical and medical grounds. Our goal now is to investigate feasible and clinically applicable treatment strategies, hence, for the purpose of the theoretical analysis, sufficiently large upper bounds on the drug administration rates are assumed. These bounds can be chosen so that they do not affect the resulting optimal treatment in practice, while ensuring that the solutions of the system remain bounded from above.

## 3 Control theory

Since the goal of drug interventions is to influence the dynamics of the system in a direction that is beneficial to the patient, considering the phenomenon from a control theory perspective is worthwhile. We introduce a new function, the control *u*(*t*), which is composed of the dosages of the chemotherapeutic drug and the antibiotics. Our goal is to bring the system to a state where *T* (*t*), i.e. the tumour size, is sufficiently low. For constant doses, this is fully answered by the results in [7], but in real-life treatments, the drug doses are typically time-dependent. In addition, the control is expected to achieve the desired state in the shortest possible time and with the fewest possible side effects. For this type of problem, the two main tasks are to construct an appropriate cost function, depending on the control and the state of the system, and to find the corresponding optimal control where the cost is minimal. Recently, optimal control methods have been increasingly applied to a number of biological ([9, 10]) and especially cancer ([11, 12, 13]) problems, making it a promising direction.

### 3.1 Theoretical background of Pontryagin’s principle

A commonly used method for finding optimal control is Pontryagin’s principle. We summarize the used notations and theorems below, further details can be found in the literature ([9, 14, 15, 16]).

Let *x ∈* ℝ^*n*^ be the function describing the state, *u ∈ U ⊂* ℝ^*m*^ be the control, where *U* is the set of admissible controls, and *f* : ℝ^*n*^ *× U →* ℝ^*n*^. Consider the equation system

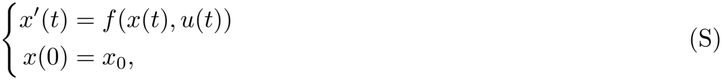

where *x*_0_ *∈* ℝ^*n*^, and the cost function

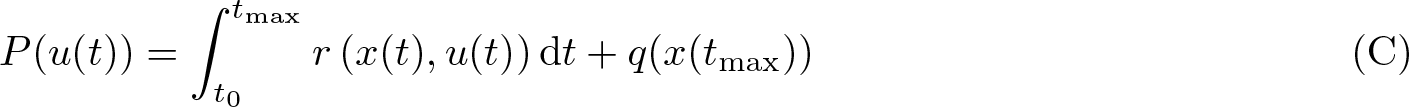

on the time interval [*t*_0_, *t*_max_], where *t*_max_ is the fixed end time. In (C), the integral term is called the running payoff, which corresponds to the harmful side effects of the control, as well as the state of the system throughout the entire treatment, while the last *q* term is the final payoff, which depends on the state of the system at the end of the treatment. We look for the optimal control *u*^*\**^(*t*) *∈ U* for which *P* (*u*(*t*)) is minimal. The corresponding optimal trajectory is *x*^*\**^(*t*), which is the solution of (S) for *u*(*t*) *≡ u*^*\**^(*t*).

Let *p* = (*p*_1_, …, *p*_*n*_) *∈* ℝ^*n*^, then we define the Hamiltonian of the system as

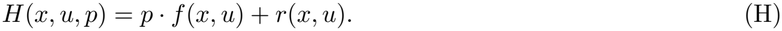

#### Theorem 3.1

(Pontryagin’s principle). *Let u*^*\**^(*t*) *be the optimal control of the problem* ((S), (C)) *and x*^*\**^(*t*) *be the corresponding optimal trajectory. Then there exists a function p*^*\**^(*t*) : [*t*_0_, *t*_*max*_] *→* ℝ^*n*^ *for which the Hamiltonian function takes its extremum in u*^*\**^, *i.e*.

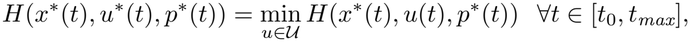

*so for every j* = 1, …, *m*

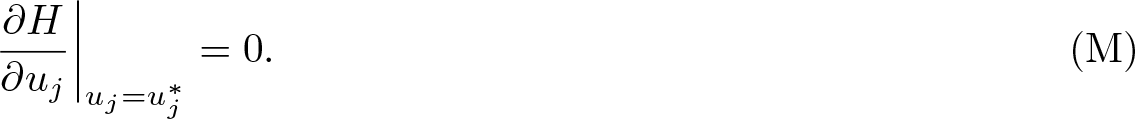

*Moreover*,

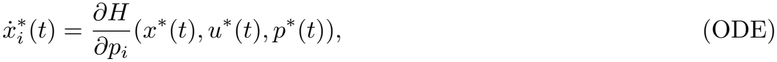

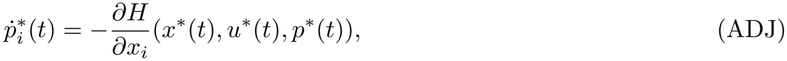

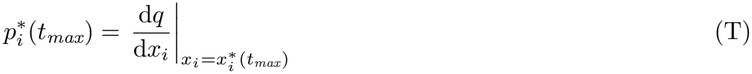

*stand for all i* = 1, …, *n, and the map t → H*(*x*^*\**^(*t*), *u*^*\**^(*t*), *p*^*\**^(*t*)) *is invariant*. □

In this case, (ADJ) is the adjoint equation and (T) is the terminal condition. The type of the extremum (minimum or maximum) is determined by the second derivative of the Hamiltonian function (H) with respect to *u*.

### 3.2 Pontryagin with a modified FBSM for the saboteur bacteria problem

Now we apply Pontryagin’s principle to the saboteur bacteria problem. The function describing the state is *x*(*t*) = (*T* (*t*), *C*(*t*), *B*(*t*), *A*(*t*))^T^, while the control is *u*(*t*) = (*γ*(*t*), *α*(*t*))^T^, thus it is composed of the doses of the chemotherapeutic drug and the antibiotics. Let the differential equation system be

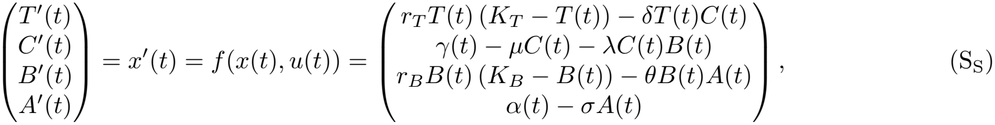

with *x*(*t*_0_) = (*T*_0_, *C*_0_, *B*_0_, *A*_0_)^T^. It is reasonable to assume that – at least in particular cases – the patient’s body cannot tolerate more than a certain amount of the drugs, so we introduce a maximum amount that can be administered, which is *γ*_max_ and *α*_max_ for chemotherapeutic drug and antibiotics, respectively. The drug dosages are obviously nonnegative quantities, so the set of admissible controls in this case is

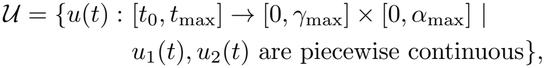

which is clearly nonempty, closed and convex in ℝ^2^. By Proposition 2.1, our model is also well-defined for piecewise continuous controls, so there exists a unique solution of the differential equation system. We aim to minimize the cost function

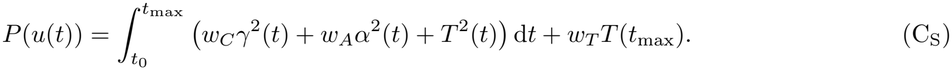

The running cost accounts for the amount of drugs and the size of the tumour throughout the treatment, and is quadratic, as commonly in the literature. Drugs administered in high doses and huge tumours are strongly toxic to the human body, which justifies the inclusion of quadratic terms in the functional. For the final payoff term, we also “penalise” the remaining tumour at the end of treatment. Since the location of the optimum is unchanged by a constant multiplier, we can assume for simplicity that one of the weights in the cost function is 1. Here, we normalize to the weight of the *T* ^2^(*t*) term in the integral and assume that the other weights (*w*_*C*_, *w*_*A*_, *w*_*T*_ ) are positive. As our goal is to eradicate the tumor, or at least reduce it as much as possible, we always choose a big *w*_*T*_ value for the cost function in the numerical simulations, to find an optimal control that results in a small residual tumor.

#### Proposition 3.2.

*There exists an optimal control u*^*\**^ *∈ U and a corresponding solution x*^*\**^ *satisfying* (S_S_), *which minimize the cost function* (C_S_).

**Proof** . The existence of such optimal control follows from standard classical results (see Chapter III of [17]), given that our controls take values from the bounded set *U* = [0, *γ*_max_] *×* [0, *α*_max_], the trajectories are uniformly bounded (see Proposition 2.1), and the running cost is given by a convex function. □

We can derive that the corresponding Hamiltonian takes the form

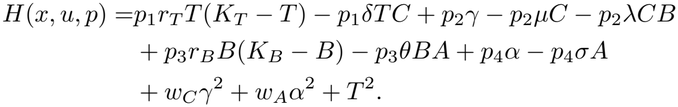

The Hessian of *H* with respect to the control *u* is

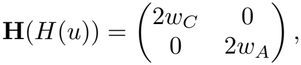

which is a positive definite matrix, so the optimal control will indeed minimize the cost function. Applying Theorem 3.1, we obtain the following equations for the optimal control and the corresponding state and adjoint variables:

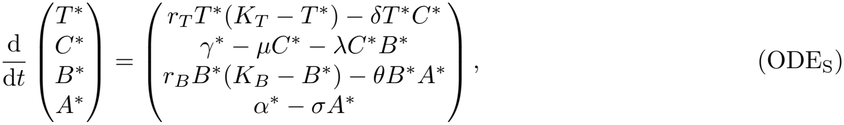

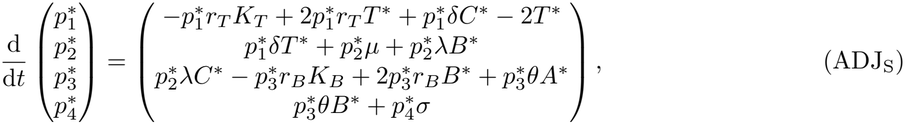

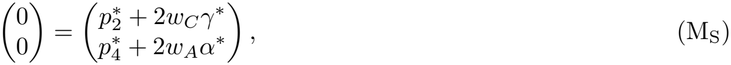

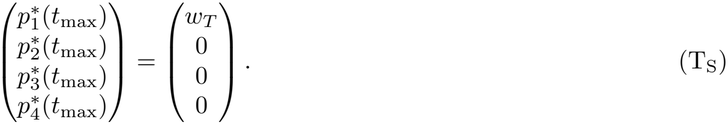

Since we need 0 *≤γ*(*t*) *≤γ*_max_ and 0*≤ α*(*t*) *≤ α*_max_ to stand for all *t∈* [*t*_0_, *t*_max_], we use this and (M_S_) to obtain the following for the control:

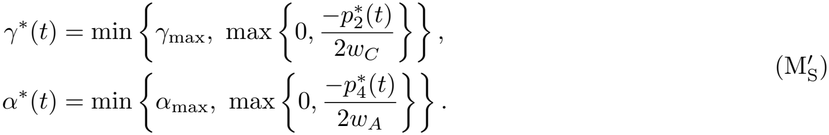

When we do not want to take the upper bounds into account, we just simply take *γ*_max_ = *∞*and/or *α*_max_ =*∞* .

The resulting system cannot be solved explicitly, so we use an iterative method: we choose constant *γ*^(0)^(*t*) *≡ γ*_0_ and *α*^(0)^(*t*) *≡ α*_0_ functions as initial estimates of the control. Using these and the initial condition *x*(*t*_0_), we can solve the subsystem (ODE_S_) forward in time. By substituting the obtained *T* ^(1)^, *C*^(1)^, *B*^(1)^ and *A*^(1)^ functions, we can solve the subsystem (ADJ_S_) backwards in time using the terminal condition (T_S_). Finally, we can express the new controls from (M_S_) using the computed 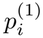 functions. Now we start the iteration: we substitute the obtained *γ*^(1)^ and *α*^(1)^ functions back into (ODE_S_) and perform the whole process again. This procedure is called the Forward-Backward Sweep Method (FBSM), was introduced by Krylov and Chernous’ko ([18]) and was first applied to biological models by Lenhart ([9]). A flowchart for the iteration is shown in Figure 1.

**Figure 1:**
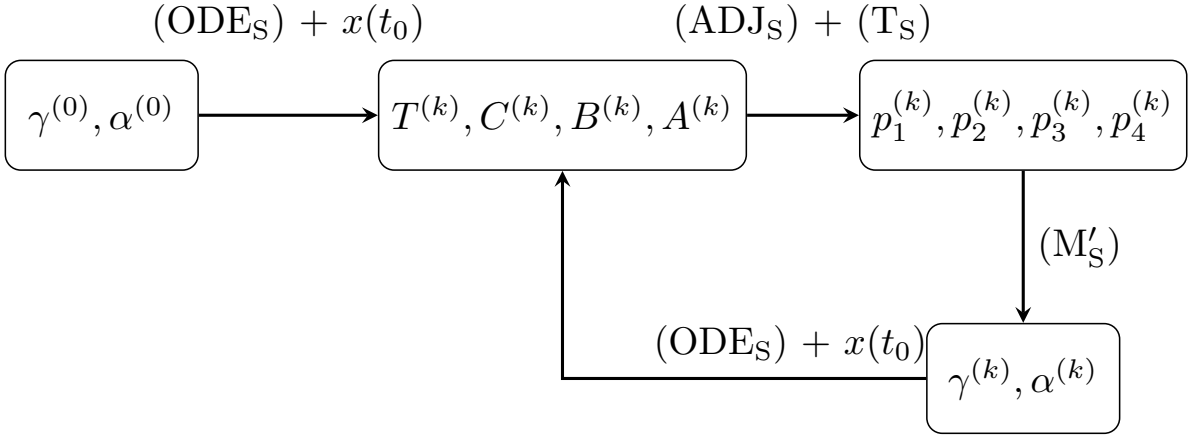
Flowchart of the FBSM. Note that for the control *u*^(*k*)^, the corresponding state response is always *x*^(*k*+1)^.

**Figure 2:**
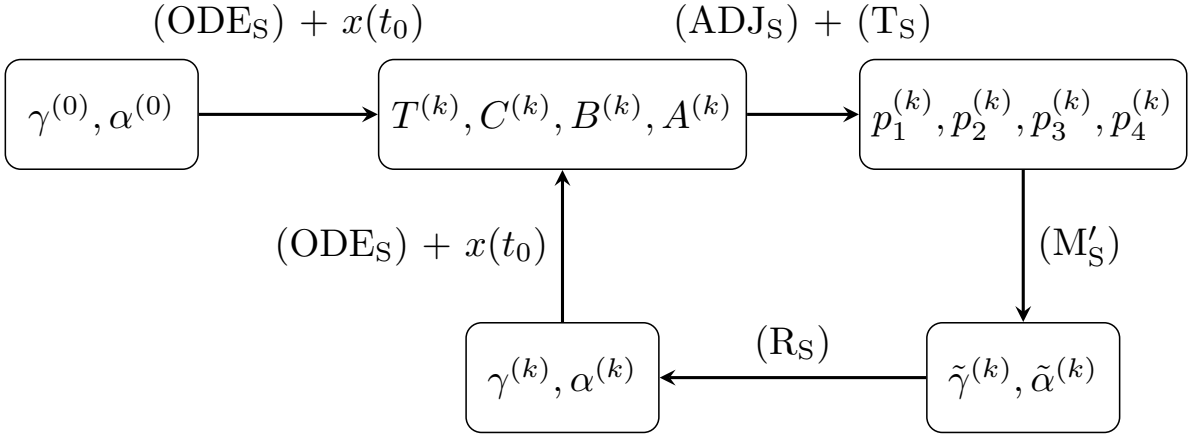
Flowchart of the modified FBSM with the relaxation parameter.

Our goal is to achieve convergence during these iterations, at which point we stop the procedure and obtain the optimal control and the corresponding solution of the system. It can be shown (see [19]) that without a terminal payoff *q* and under some reasonable assumptions, the FBSM is indeed convergent. Since in our case the boundedness of the controls is important, we use (M^*′*^_S_) instead of (M_S_) in the iteration. We can do this without affecting the conditions for convergence ([19], Remark 3.6.2). Although this form of the FBSM is sufficient for solving certain problems with terminal conditions ([20], Example 7.2.), the procedure does not converge in all cases. Therefore, we modify the method based on [21] and introduce a relaxation parameter *ω ∈* [0, 1). Now, at each step, when we update the control, we take the convex combination of the last computed control and the previous one at the reset. That is, if we have *u*^(*k−*1)^ after the (*k −*1)th iteration, and we compute *ũ*^(*k*)^ during the next one, then we take

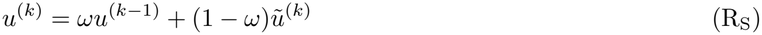

and substitute that back into (ODE_S_). In this way, we can achieve convergence in cases where we could not before. For now, we use fixed *ω* values, but other methods to facilitate and accelerate convergence are mentioned in Section 4.2.

It is important to note that for every iteration step during the FBSM, we always get controls that are in *U* . The constant functions used as initial estimators are obviously piecewise continuous, while in (M^*′*^_S_), the *ũ*^(*k*)^’s are expressed in terms of constant bounds and *p*^(*k*)^’s, the latter being solutions of a differential equation system and thus continuously differentiable. And since *U* is convex, the relaxation step also does not lead out of it.

The iteration is stopped when the difference between consecutive controls is small enough, which we measure with the widely used criterion

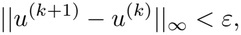

where *ε* is given. Although convergence of both the state and adjoint variables is also essential, the continuous dependence of the state and adjoint solutions on the control allows us to assess convergence by monitoring changes in the control alone.

## 4 Numerical results

### 4.1 Simulations

We ran numerous simulations for different scenarios and parameter combinations. In every case, we had the following assumptions: the effect of the chemotherapeutic drug is more harmful than the effect of the antibiotics, so we always take *w*_*C*_ *> w*_*A*_; and both drugs are at most as destructive as the tumour burden, i.e., *w*_*C*_, *w*_*A*_ *≤* 1.

Since we want as little amount of the tumour as possible by the end of the treatment, *w*_*T*_ is significantly bigger than the other weights in the cost function (C_S_). Finally, we always take *γ*^(0)^ = *α*^(0)^ = 0 for the initial guesses ([9] says that 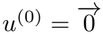 is almost always sufficient). The time interval for every simulation is [*t*_0_, *t*_max_] = [0, 10], and the constant in the convergence stopping criterion is *ε* = 10^*−*4^. In order to draw more reasonable conclusions, most of the parameters are fixed and only a few are changed between simulations plotted in the same figure. The fixed parameters can be found in Table 2. We used the relaxation constant *ω* = 0.88 for all of the simulations . Other choices for *u*^(0)^ and *ω* are mentioned in Section 4.2.

**Table 2:**
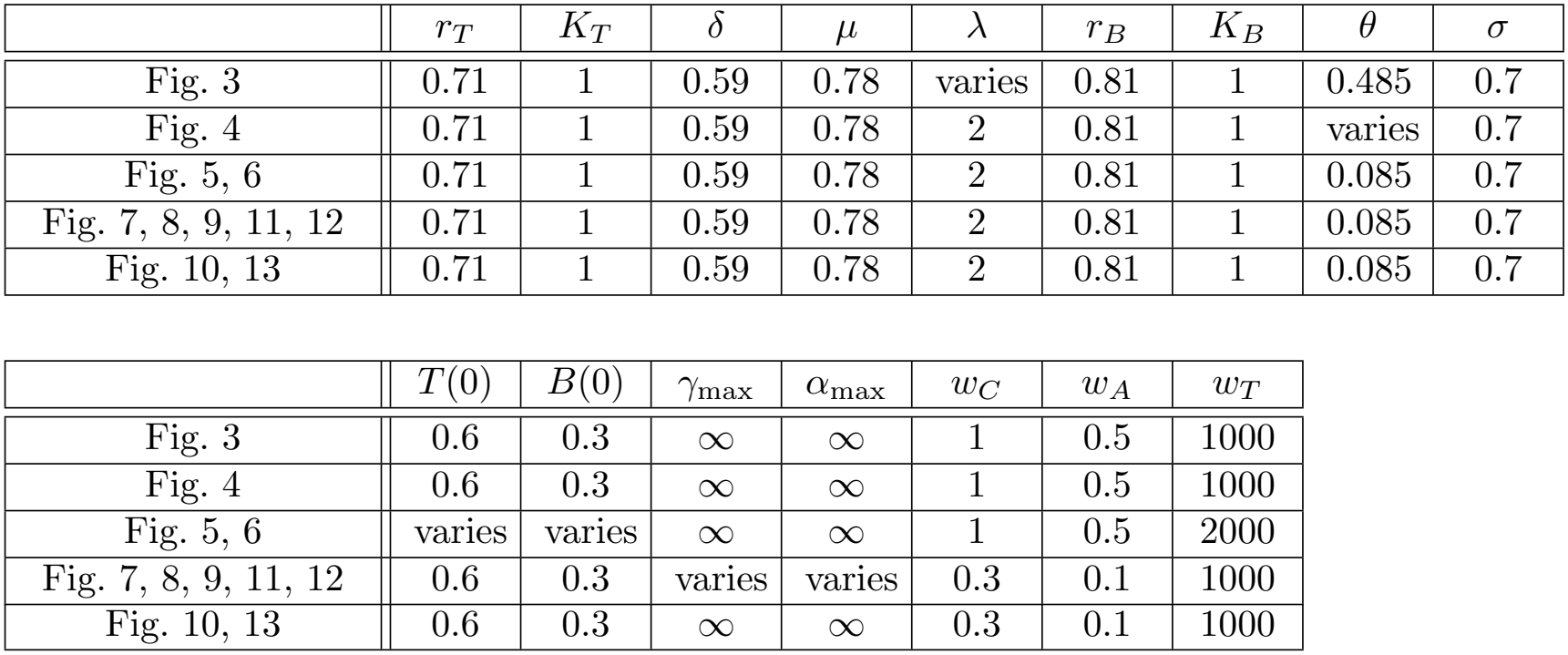
Fixed parameters in the simulations.

|  | *r*<sub>*T*</sub> | *K*<sub>*T*</sub> | δ | μ | λ | *r*<sub>*B*</sub> | *K*<sub>*B*</sub> | θ | σ |
|---|---|---|---|---|---|---|---|---|---|
| Fig. 3 | 0.71 | 1 | 0.59 | 0.78 | varies | 0.81 | 1 | 0.485 | 0.7 |
| Fig. 4 | 0.71 | 1 | 0.59 | 0.78 | 2 | 0.81 | 1 | varies | 0.7 |
| Fig. 5, 6 | 0.71 | 1 | 0.59 | 0.78 | 2 | 0.81 | 1 | 0.085 | 0.7 |
| Fig. 7, 8, 9, 11, 12 | 0.71 | 1 | 0.59 | 0.78 | 2 | 0.81 | 1 | 0.085 | 0.7 |
| Fig. 10, 13 | 0.71 | 1 | 0.59 | 0.78 | 2 | 0.81 | 1 | 0.085 | 0.7 |
|  | *T*(0) | *B*(0) | γ<sub>max</sub> | α<sub>max</sub> | *w*<sub>*C*</sub> | *w*<sub>*A*</sub> | *w*<sub>*T*</sub> |  |  |
| Fig. 3 | 0.6 | 0.3 | ∞ | ∞ | 1 | 0.5 | 1000 |  |  |
| Fig. 4 | 0.6 | 0.3 | ∞ | ∞ | 1 | 0.5 | 1000 |  |  |
| Fig. 5, 6 | varies | varies | ∞ | ∞ | 1 | 0.5 | 2000 |  |  |
| Fig. 7, 8, 9, 11, 12 | 0.6 | 0.3 | varies | varies | 0.3 | 0.1 | 1000 |  |  |
| Fig. 10, 13 | 0.6 | 0.3 | ∞ | ∞ | 0.3 | 0.1 | 1000 |  |  |

The simulations were performed with the software Wolfram Mathematica (version 14). In Section 4.1, there are two images for every simulation: the one on the left shows the drug doses (chemotherapy with solid, antibiotics with dashed), while the one on the right shows the size of the tumour (with solid) and the bacteria population (with dashed) over the course of the therapy using the drug dosages shown on the left. The related graphs have the same colour in both images.

Based on these simulations, we can see what the general treatment process is. First, we give a larger amount of antibiotics, while trying to “keep the tumour in check” with a small amount of chemotherapy. When we have successfully reduced the bacterial population - and with it the strength of the sabotaging effect - then we increase the amount of the chemotherapeutic drug to eradicate the tumour. As the tumour size decreases towards the end of the treatment, we can reduce the amount of the chemotherapy too.

#### 4.1.1 Sabotaging effect

First we investigate how the strength of the bacteria’s sabotaging effect influences the needed therapy. This means that we consider simulations with different *λ* values. Keep in mind that larger *λ* means stronger sabotaging effect. These simulations are shown in Figure 3 which confirms what we expect. If the sabotaging effect is stronger, then it is more important to get rid of the bacteria, so we need a larger amount of the antibiotics. We can also see that if *λ* is large, then we need to eradicate the bacteria completely, while for a smaller *λ*, it is enough to moderate the size of the population.

**Figure 3:**
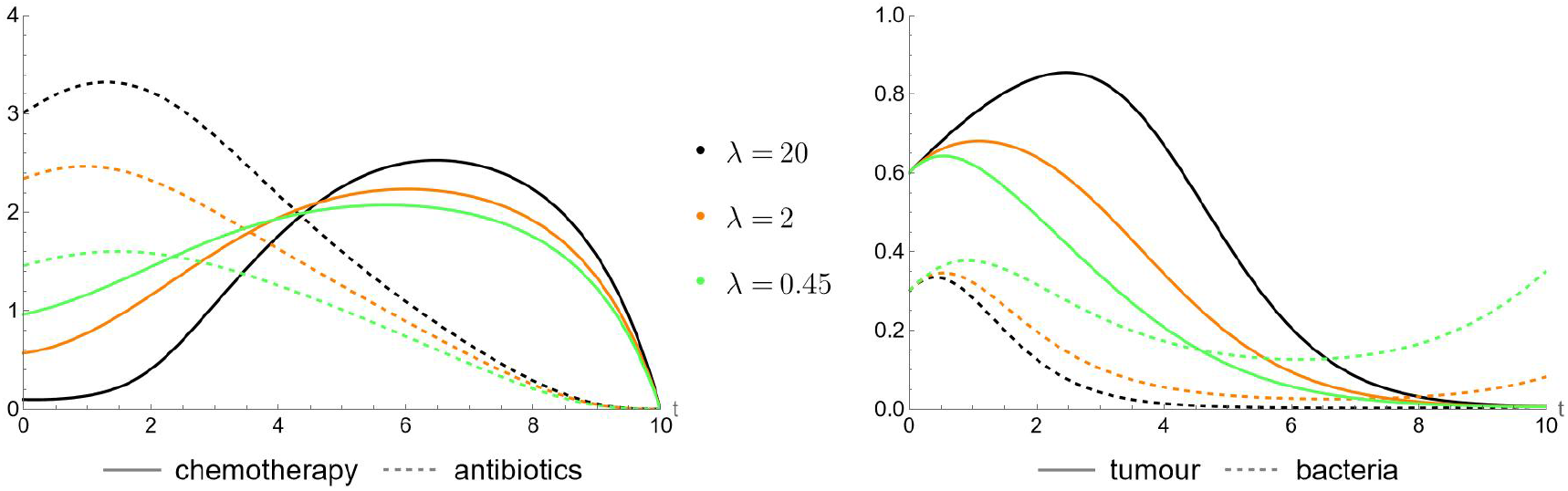
The influence of the strength of the sabotaging effect. Fixed parameters are listed in Table 2.

#### 4.1.2 Effectiveness of the antibiotics

Next, we investigate the impact of the antibiotics’ effectiveness on the treatment. Larger *θ* means that the antibiotics is more effective, i.e., the bactericidal activity is stronger. These simulations are depicted in Figure 4. We can see that if the antibiotics is more powerful, then it is enough to use a smaller amount of it. Also if it is not really effective, then we are not able to kill a substantial part of the bacterium population, and we have to compensate it with a larger amount of chemotherapeutic drug.

**Figure 4:**
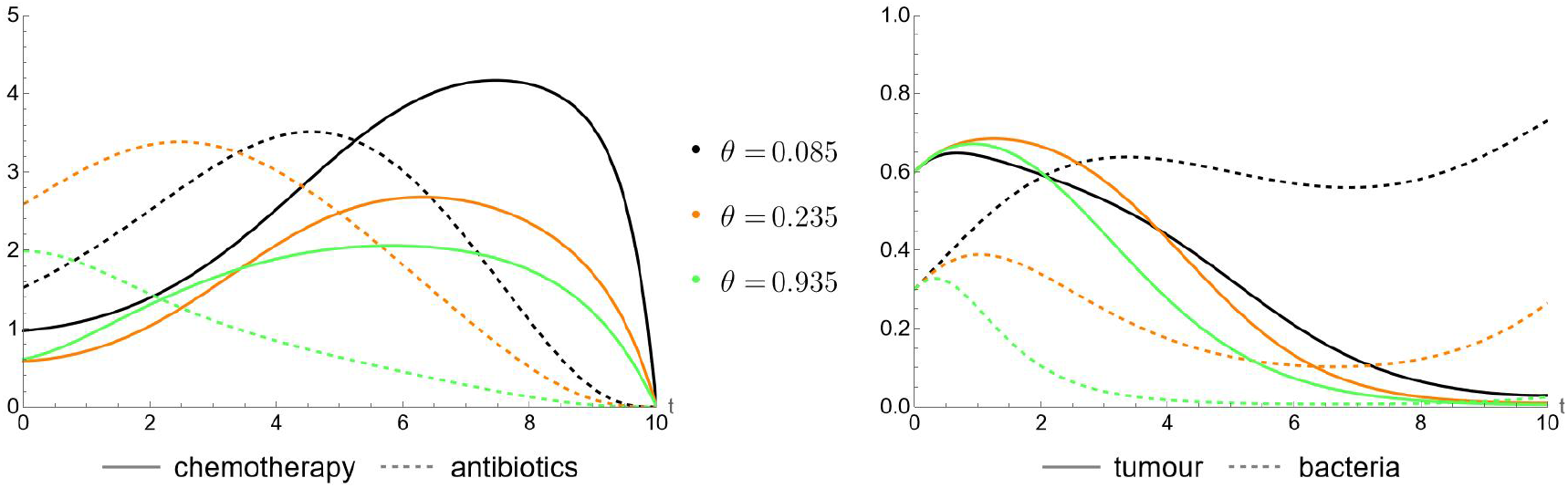
Effectiveness of the antibiotics. Fixed parameters are listed in Table 2.

#### 4.1.3 Initial values

Since treatments may differ depending on the size of the tumour and the bacterial population when therapy is started, simulations were created with different *T* (0) and *B*(0) initial values. We can see in Figure 5 that the optimal outcome is achievable in every scenario, but the therapies are indeed slightly different. Figure 6 shows the same simulations as Figure 5 for more transparency.

**Figure 5:**
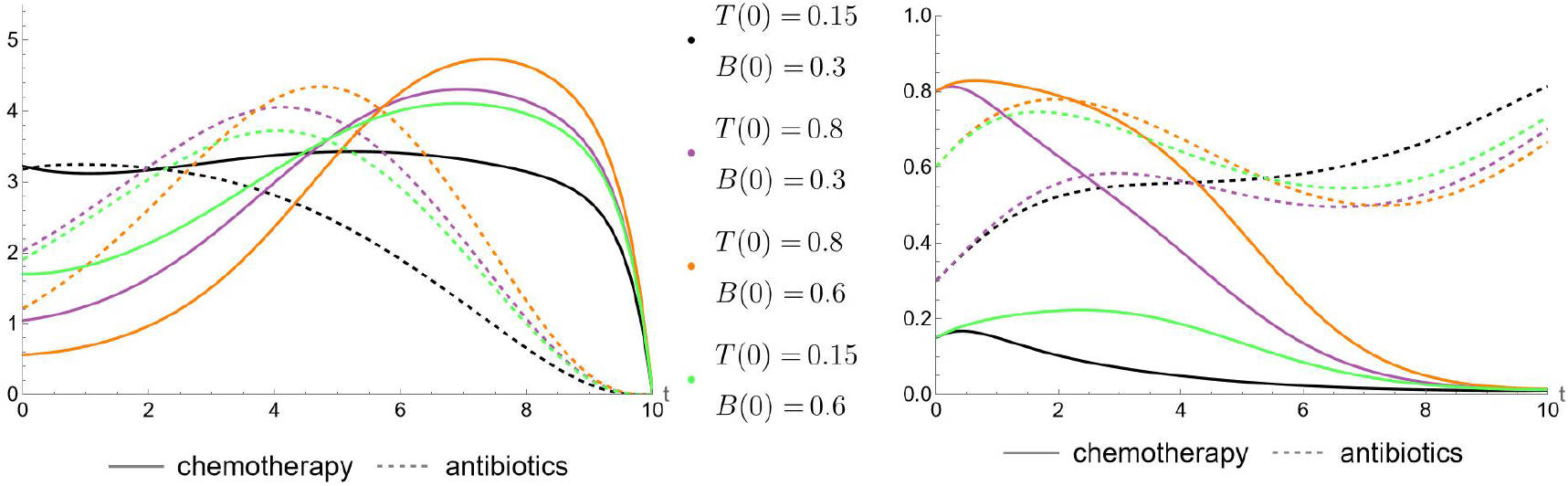
Simulations with different initial values. Fixed parameters are listed in Table 2.

**Figure 6:**
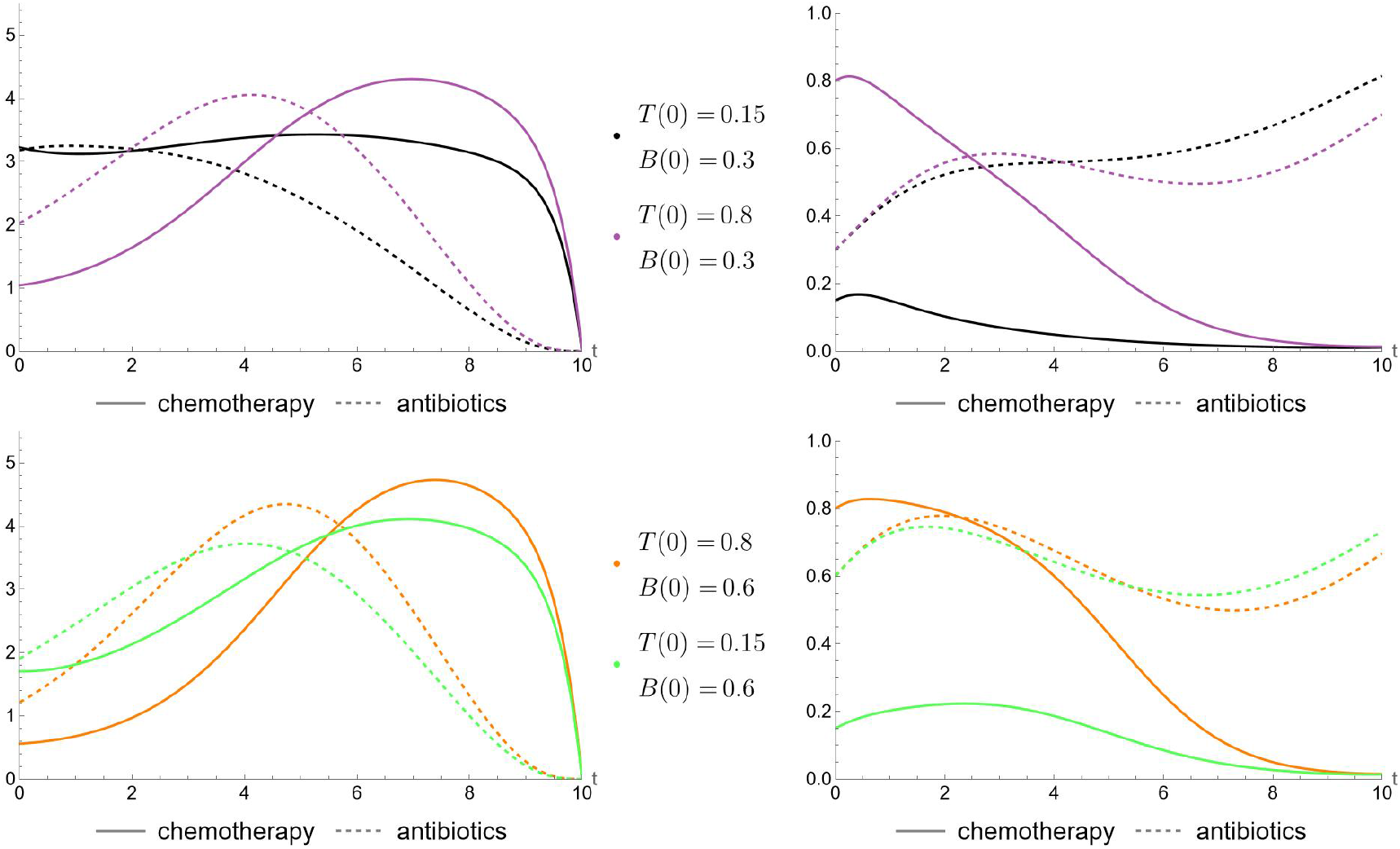
Simulations with different initial values. Fixed parameters are listed in Table 2.

#### 4.1.4 Controls with bounds

As mentioned earlier, we should also consider scenarios where the maximum concentration of the drugs is limited. The optimal result is still achievable in some of these situations, the corresponding simulations are depicted in Figure 7. For easier comparison of the drug doses, Figure 8 shows the same simulations as Figure 7, but only two at a time: the case without upper bounds (always with the darker colour), and a case where we have different kind if upper bounds for either the chemotherapy, the antibiotics or both (with lighter colour). We can see if one of the drugs has an upper bound, then we need to start giving a higher concentration of it earlier, and in some cases increase the amount of the other drug too.

**Figure 7:**
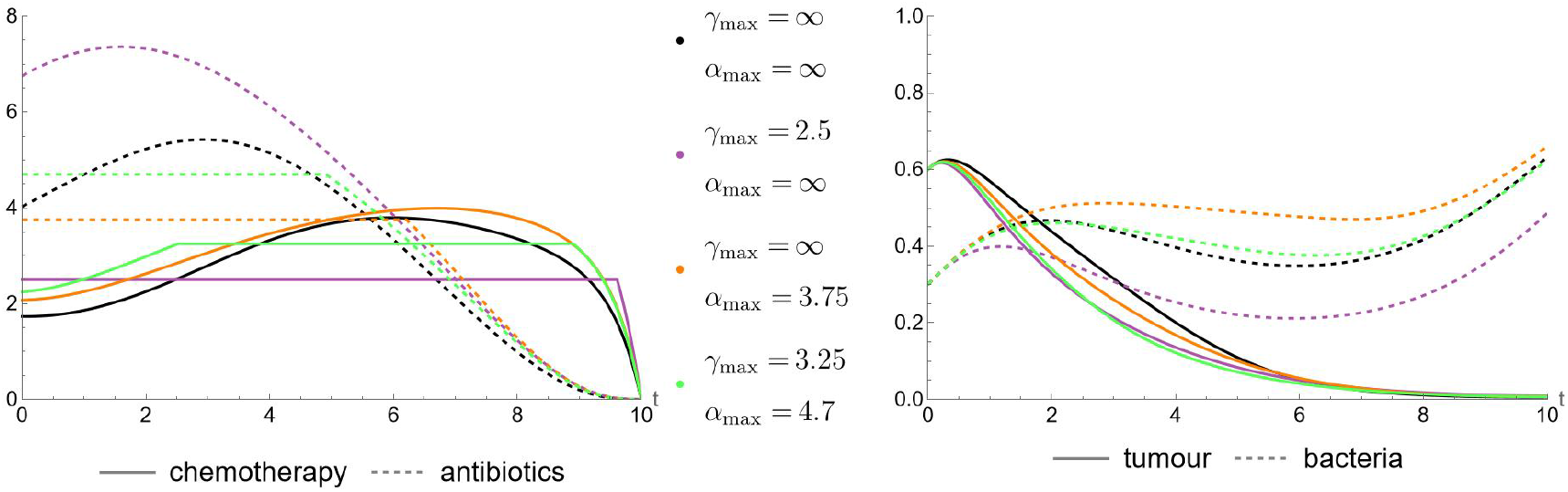
Simulations of successful therapies with different upper bounds for the controls. Fixed parameters are listed in Table 2.

**Figure 8:**
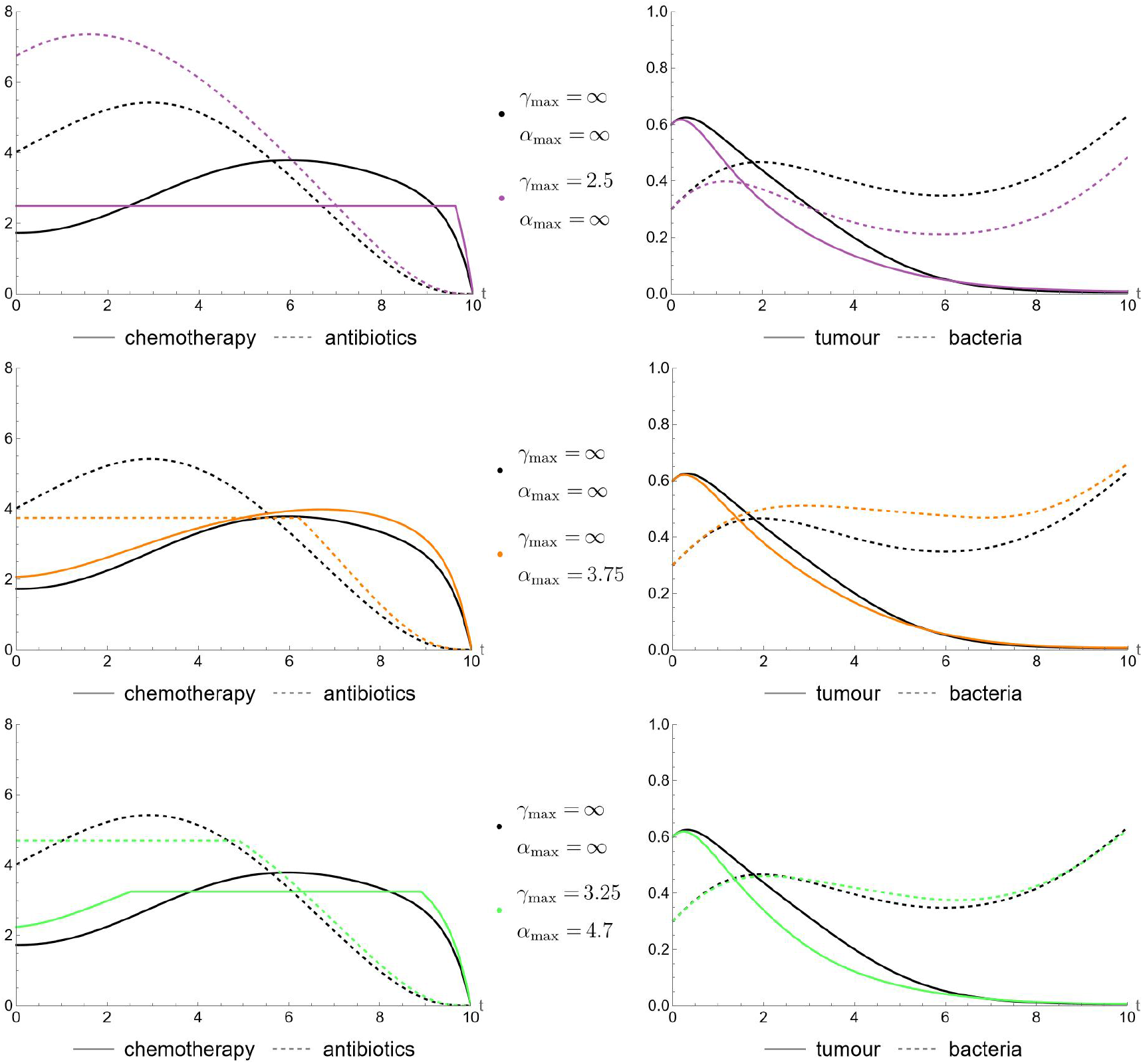
Simulations of successful therapies with different upper bounds for the controls. Fixed parameters are listed in Table 2.

##### Remark 4.1

(Tumour persistence). Obviously, it can happen that the upper bounds are too low and the therapy fails even with the optimal control. Although a complete characterization of the cases in which tumour eradication fails is beyond the scope of this work, we can identify a simple sufficient condition under which tumour eradication by the end of the treatment interval is impossible for any admissible therapy. Suppose that the upper bound on the chemotherapy control, *γ*_max_, satisfies *K*_*T*_ *µr*_*T*_ */δ > γ*_max_. By comparing this condition with Theorems 2.3 and 2.7 in [7], we obtain *F*_2_ *>* 0, corresponding to scenarios in which the tumour cannot be eradicated under the constant chemotherapy regimen *γ*(*t*) *≡ γ*_max_. Since *γ*_max_ represents the maximum admissible chemotherapy dose, no smaller chemotherapy dose can achieve tumour eradication either. Moreover, because ℱ_2_ is independent of the antibiotic dose, varying the antibiotic administration over time cannot alter this conclusion. Consequently, under the above condition, tumour persistence is guaranteed for every admissible choice of the treatment controls. We do not consider here the problem of minimizing the residual tumour burden in this case. The purpose of this Remark is solely to identify a simple sufficient condition under which tumour eradication is impossible, irrespective of the admissible treatment strategy.

Figure 9 shows a simulation in which the bound for the chemotherapeutic drug is so low that we cannot compensate for it even with a really high dose of antibiotics and we cannot perform a successful therapy.

**Figure 9:**
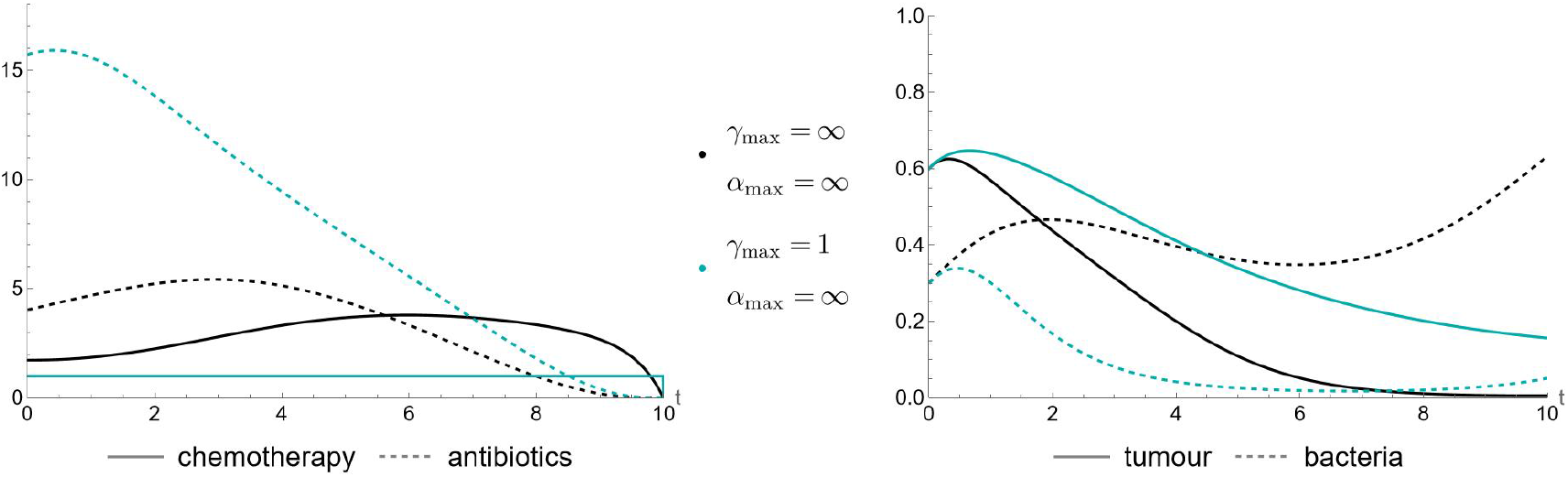
Simulation with upper bounds for the controls where the therapy does not succeed. Fixed parameters are listed in Table 2.

#### 4.1.5 Drug cost vs. effectiveness

Another interesting direction for future research would be to investigate how the optimal treatment strategies differ between chemotherapeutic agents with distinct pharmacological properties. Consider, for instance, two drugs: one with a high tumour-cell killing rate (i.e., a large value of *δ*), but also a high toxicity towards healthy cells, reflected by a large value of *w*_*C*_, and another with a lower killing rate and correspondingly lower toxicity. Which drug would be preferable, and which would result in a lower overall treatment cost? It is conceivable that the additional tumour-killing effect of the more potent drug may not justify its use if the associated burden becomes too high.

Unfortunately, our model contains a relatively large number of parameters, which complicates a purely theoretical comparison of different drugs. Each of the two drugs is characterized by three parameters: the killing rate, the absorption rate, and its contribution to the cost functional through the corresponding burden coefficient. In addition, the system contains several other parameters that can substantially influence the resulting optimal treatment, as demonstrated in the preceding analysis and simulations. Consequently, a meaningful comparison of different drugs and the derivation of reliable conclusions would require specific experimental data and parameter estimates, using hypothetical parameters may lead to false deductions.

Nevertheless, we would like to emphasize a potentially valuable application of the optimal control framework. For each admissible combination of drug parameters, an optimal control problem can be formulated and an optimal treatment can be obtained. Different drugs give rise to different parameter combinations and, consequently, potentially different optimal treatment strategies and associated cost functional values. Comparing these values would provide a basis for identifying the most favourable treatment among the available alternatives. This illustrates a potential practical benefit of optimal control, even when the resulting control functions cannot be implemented directly in clinical practice. Although the computed continuous dosing profiles may be difficult to translate into feasible treatment protocols, comparisons between candidate drugs could still provide valuable guidance for drug selection. In particular, the framework could help identify which agents are most promising for further investigation, thereby supporting the medical decision-making process rather than replacing clinical judgement.

### 4.2 Convergence analysis

The success of the FBSM relies on the convergence of its iterations, making convergence a central issue in determining the effectiveness and applicability of the method. As mentioned earlier, convergence can be guaranteed for certain problems or under specific conditions, but this is not always the case. Although establishing convergence is beyond the scope of the present study, we briefly discuss several relevant approaches that may be of interest to readers wishing to explore this topic further. Figure 10 illustrates a convergent iteration, where every third iterate of the control functions *γ*(*t*), *α*(*t*) and the corresponding states *T* (*t*) and *C*(*t*) are plotted.

**Figure 10:**
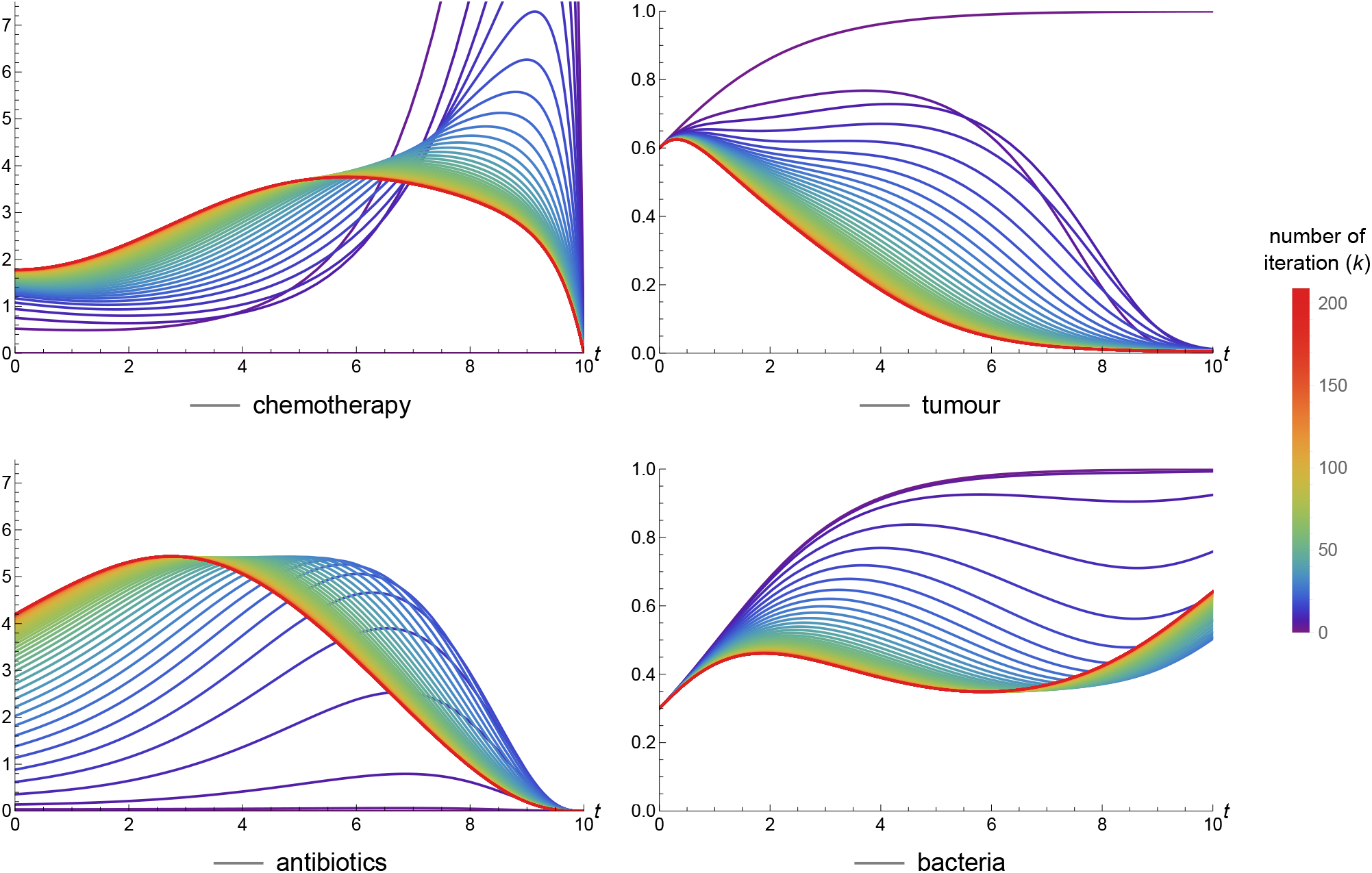
Convergence of controls and state variables. The colours of the graphs correspond to the iteration number of the given control/state, as shown in the bar legend, with the initial control guesses being dark purple, and the optimal trajectories achieved after the convergence are red. Corresponding parameters are listed in Table 2. Every third iterate is plotted.

#### 4.2.1 Relaxation methods

Relaxation modifies the convergence behaviour of the iteration without changing its fixed points, thereby providing flexibility in the choice of the relaxation parameter *ω* in (R_S_). Since *ω* determines the relative weight assigned to the control obtained in the previous iteration, smaller values of *ω* generally allow for faster updates, whereas larger values may improve the stability and convergence properties of the iteration. Consequently, choosing an appropriate relaxation parameter involves a trade-off between convergence speed and robustness. Figure 11 illustrates how *N*, the number of iterations needed for convergence heavily depends on the value of *ω*.

**Figure 11:**
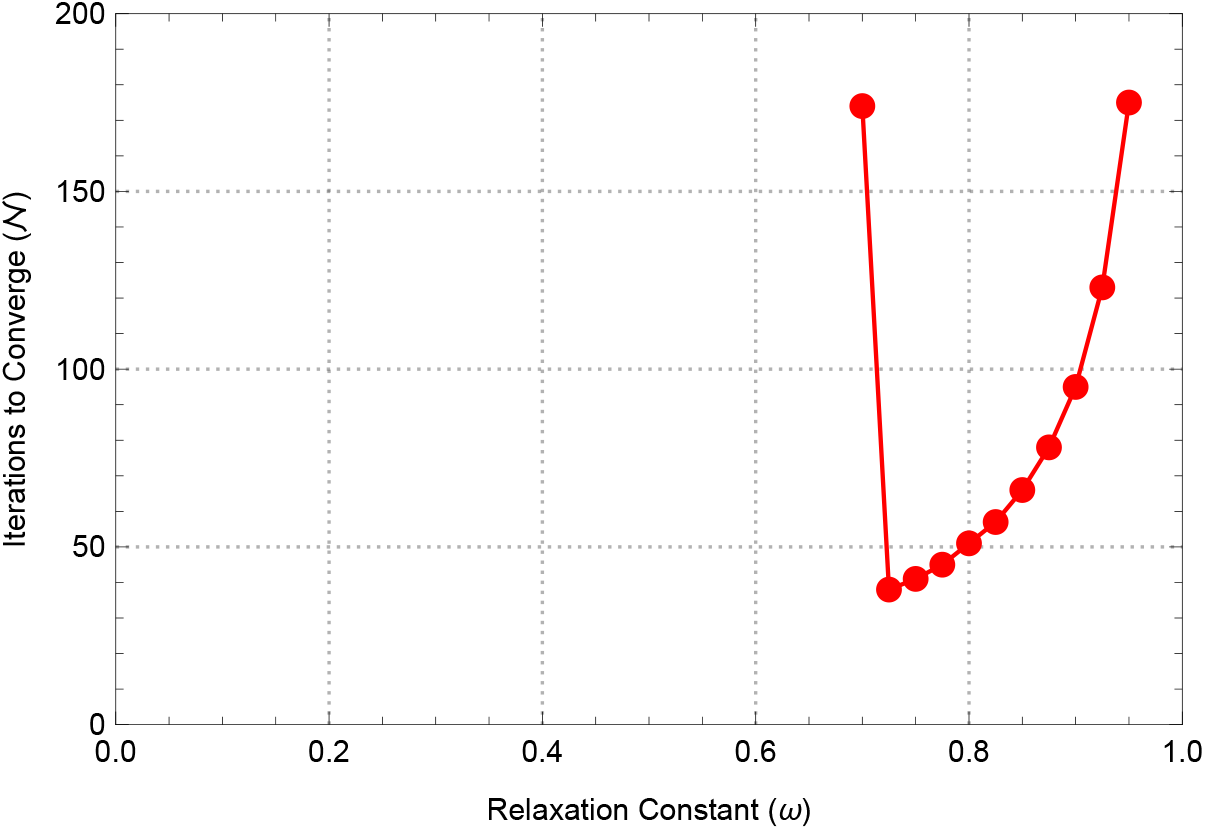
*N* against *ω ∈* [0, 1) in increments of 0.025. For a given *ω* value without a corresponding marker there is no convergence within 200 iterations. The rest of the parameters are listed in Table 2, with *γ*_max_ = 2.5, *α*_max_ = *∞*.

Several approaches can be considered to accelerate convergence. In the description of our FBSM algorithm and throughout the numerical simulations presented in this work, we use a fixed relaxation parameter. However, *ω* can also be adjusted dynamically during the iteration. Although adaptive relaxation strategies have been discussed in the literature ([9, 21]), their practical implementation is not yet standardized, and the choice of an appropriate indicator for updating *ω*, as well as the update rule itself, requires further consideration. Possible approaches include gradient-descent-type strategies and adaptive updates based on changes in the distances between successive control iterates. Investigating such strategies, however, is beyond the scope of the present work.

An alternative approach is to formulate the FBSM as a fixed-point iteration, as described in [21]. This perspective opens the door to applying established fixed-point acceleration techniques, thereby modifying the iteration itself rather than merely adjusting a fixed relaxation parameter. Nevertheless, the convergence analysis becomes more involved for systems with multiple state and control variables, such as the model considered in this study. As explicitly noted in [21], the Wegstein method may encounter difficulties in multidimensional settings.

#### 4.2.2 Different initial guesses for the controls

As the initial control in all of our simulations, we used *u*^(0)^ *≡*0, as is commonly done in the literature. It is important to note, however, that alternative initial guesses may lead to faster convergence. In particular, when the controls are bounded both from below and above, it is also natural to consider, as constant initial guesses, the choices *u*^(0)^ *≡ u*_max_ and *u*^(0)^ *≡ u*_max_*/*2.

Based on the simulations presented above, we observed that the optimal therapy typically consists of a higher antibiotic dose combined with a lower chemotherapy dose during the initial phase of treatment, followed by a decrease in the antibiotic dose and an increase in the chemotherapy dose during the latter part of the treatment interval. This suggests a natural starting point if we move beyond constant initial controls. Specifically, for the antibiotic control, we take the maximal dose during the first half of the treatment interval and zero during the second half, whereas for the chemotherapy control we use the opposite pattern: zero during the first half and the maximal dose during the second half. If no finite upper bound is imposed on the controls, the maximal value is replaced by a suitably chosen positive constant. It is important to note that these initial guesses also belong to *U*.

Let us take the case with the parameters listed in Table 2 for Figure 12 with *γ*_max_ = 3.25, *α*_max_ = 4.7 and *ω* = 0.88. Then with the initial control guesses *u*^(0)^ *≡* 0, *u*^(0)^ *≡ u*_max_*/*2 and *u*^(0)^ *≡ u*_max_ we have convergence in *N* = 94, *N* = 88 and *N* = 69 iterations, respectively. If we use the “half and half” initial guess suggested above, convergence occurs after *N* = 83 iterations. This is illustrated in Figure 12.

**Figure 12:**
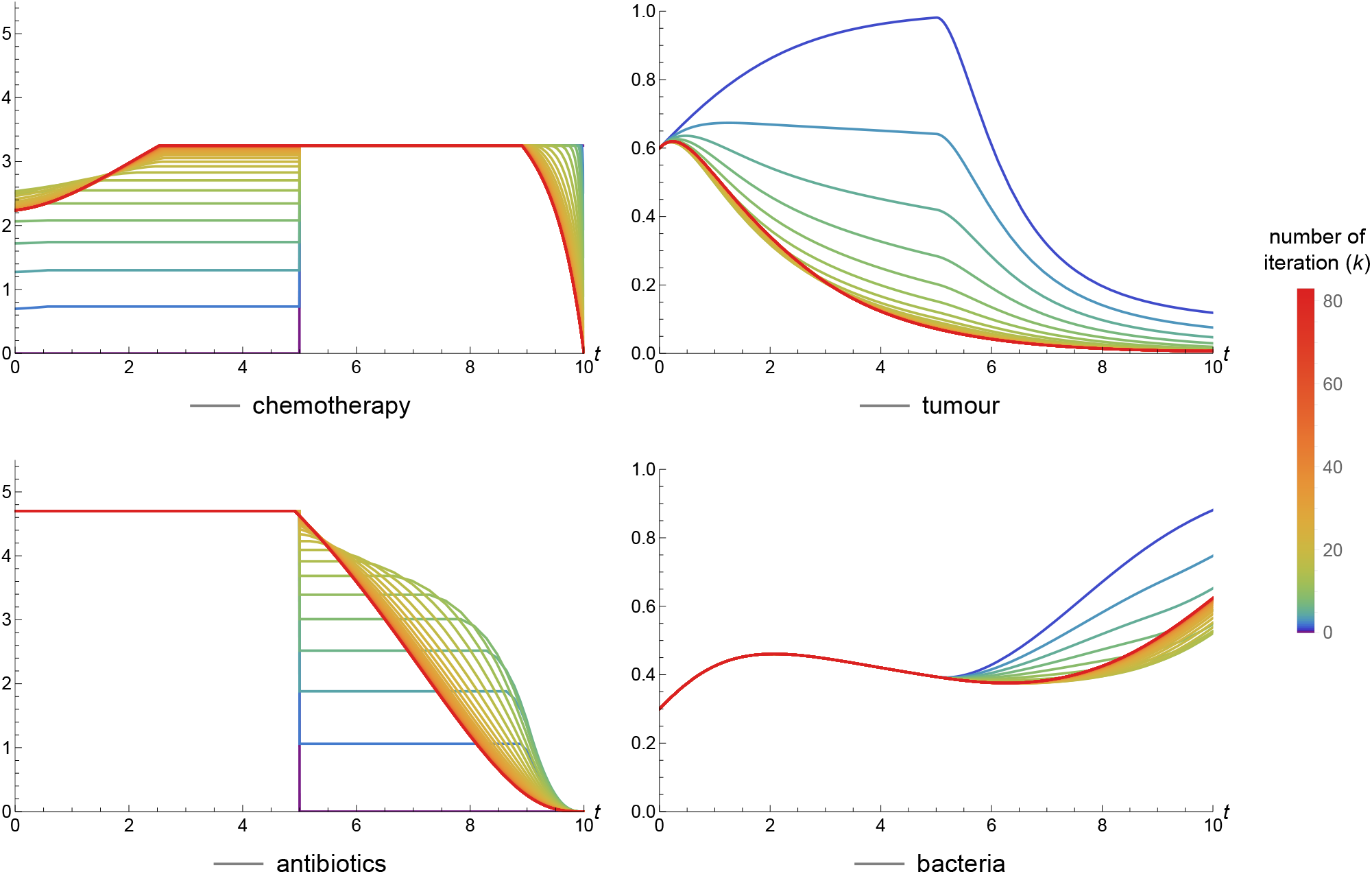
Convergence of controls and state variables from the “half and half” initial control guess. The colours of the graphs correspond to the iteration number of the given control/state, as shown in the bar legend, with the initial control guesses being dark blue, and the optimal trajectories achieved after the convergence are red. Parameters are listed in Table 2, with *γ*_max_ = 3.25, *α*_max_ = 4.7. Every second iterate is plotted.

### 4.3 Discrete controls

Although continuous drug administration is possible in some cases, like in vitro experiments or IV therapies, we investigate the effect of an impulsive (discrete in time) drug therapy. For this, we use a modified, impulsive version of the differential equation model, like in Sections 3.2 or 4.4 in [7]. The idea is to compute the continuous drug treatment – i.e. the optimal control –, then construct a discretized control based on that. After that, using simulations, we compare the effects of the continuous and the impulsive therapies.

We divide the time interval [0, *t*_max_] into *d* identical subintervals:

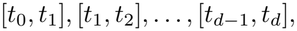

where 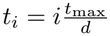 for every *i* = 0, 1, …, *d*. During the course of an impulsive therapy, we want to give approximately the same amount of drugs as we would using the continuous optimal functions *γ*(*t*) and *α*(*t*). So we define the discrete doses as

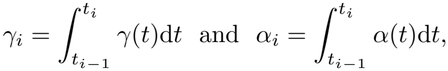

where *γ*(*t*) and *α*(*t*) are the continuous optimal controls and *i* = 1, …, *d*. This way the sum of the *γ*_*i*_ values will be “close” to the integral of *γ*(*t*) over [0, *t*_max_], if *d* is “large enough”. In particular, if we assume a constant *γ*(*t*) *≡ γ* function, then we have equality, since

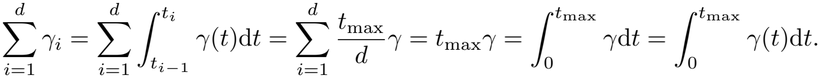

This similarly stands for the *α*_*i*_ values. We administer the *γ*_*i*_ and *α*_*i*_ doses at times 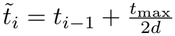, i.e. the midpoint of the interval [*t*_*i−*1_, *t*_*i*_], for every *i* = 1, …, *d*.

A simulation illustrating the effect of impulsive therapy is shown in Figure 13. The figure contains three subfigures: on the top, (a) shows the drug doses (*γ*(*t*), *α*(*t*)), where the *y*-axis on the left belongs to the continuous doses, while the *y*-axis on the right belongs to the impulsive doses. In the bottom row, (b) shows the drug concentrations (*C*(*t*), *A*(*t*)) during the treatment, while (c) shows the tumour and bacteria population sizes (*T* (*t*), *B*(*t*)), and the continuous and dashed lines correspond to the continuous and impulsive drug administrations, respectively. In every subfigure, the *x*-axis is for time, with [*t*_0_, *t*_max_] = [0, 10]. For these simulations the parameters are listed in Table 2, and the number of impulsive doses is *d* = 12. Running simulations with different number of impulsive doses shows that increasing the number of doses leads to a closer estimation of the effect of the continuous drugs, as expected.

**Figure 13:**
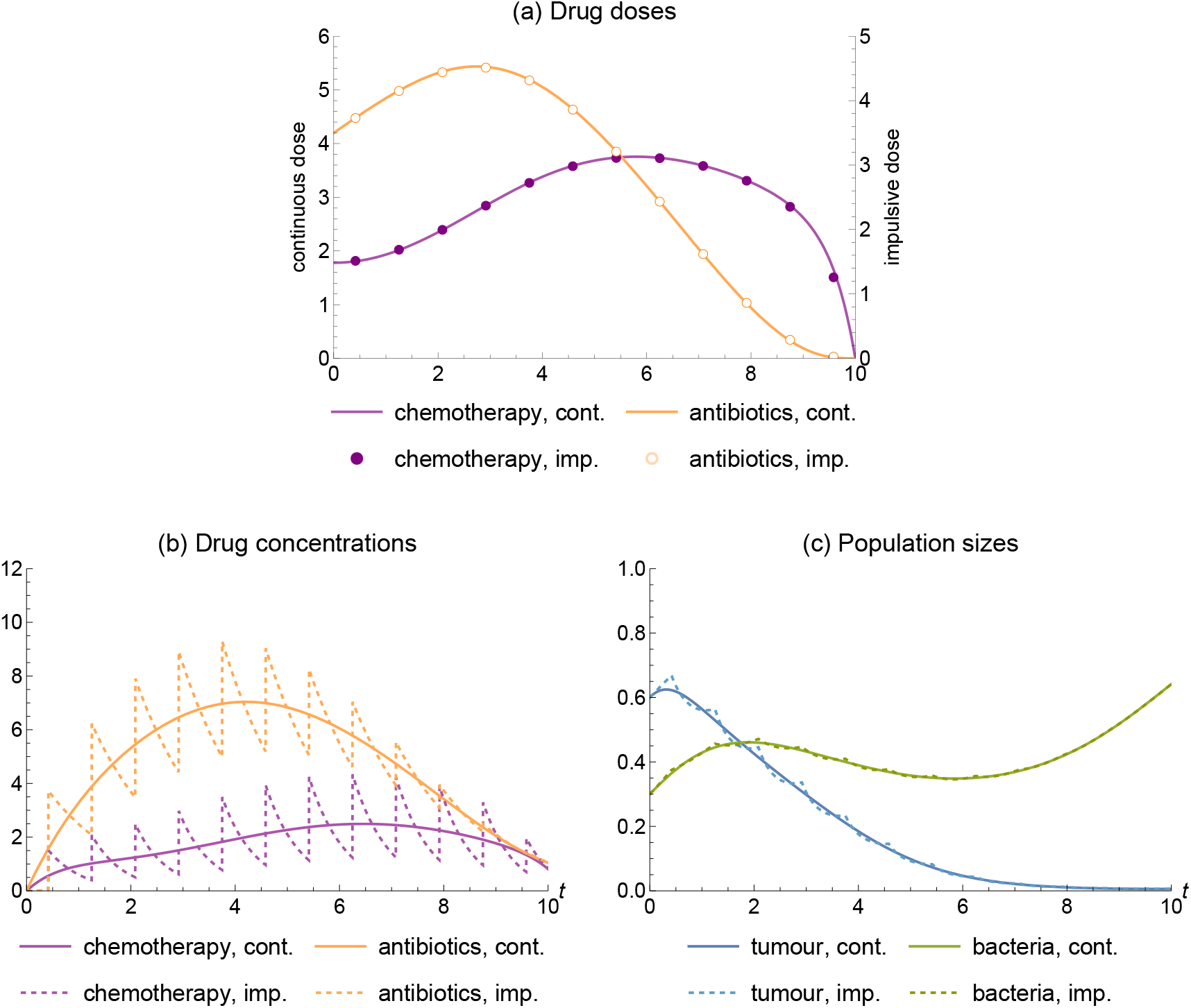
Comparison of the effects of a continuous and the relating impulsive drug therapy with *d* = 12 drug doses over the time course [0, 10]. Continuous and dashed lines correspond to the continuous and the impulsive therapies, respectively. Parameters are listed in Table 2.

## 5 Conclusions

In this work, we extended a recently developed mathematical model capturing the complex interplay between cancer dynamics, chemotherapeutic treatment, and intratumour bacteria capable of drug metabolism. By introducing time varying antibiotic co-therapy as a complementary control variable to the chemotherapeutic drug, we demonstrated that multi-drug scheduling has the potential to mitigate microbial interference and improve overall therapeutic efficacy. Our numerical investigations, based on Pontryagin’s principle, and numerically stabilized by a relaxation method within the forward-backward sweep algorithm, highlighted how optimal treatment regimes shift across various parameter regions. Furthermore, by evaluating an impulsive discrete-dose formulation across multiple time scales, we investigated realistic, scheduled clinical drug administration regimes.

As general conclusion, we may say that the optimal therapy typically consists of a higher antibiotic dose combined with a lower chemotherapy dose during the initial phase of treatment, followed by a decrease in the antibiotic dose and an increase in the chemotherapy dose during the latter part of the treatment interval. This feature of the optimal control is enhanced if the sabotaging effect of the bacteria is stronger, their initial density is higher, or the antiobiotics is less effective. We found similar conclusions when considering impulsive, discrete-dose scheduling of the drugs, resembling more real life treatment regimens. We also investigated how the rate of convergence to optimal control is depending on the initial guess in FBSM, and on the relaxation constant.

Naturally, our model has a number of limitations, such as mass action terms rather than other type of dose-response, the assumption of a well-mixed bacteria-tumour system, and the weights in the cost functions are not based on hard quantitative data, only biological intuitions.

While we employ the fixed-time, free-endpoint formulation of Pontryagin’s maximum principle, as detailed in Section 3.1, an alternative approach is to formulate the problem as a free-time, fixed-endpoint optimal control problem. In this setting, the treatment duration is not prescribed in advance; instead, the objective is to reach a prescribed low tumour burden by the end of treatment. This formulation is particularly relevant in our model, since the tumour population may converge to zero asymptotically but cannot reach zero in finite time. Furthermore, the treatment duration itself can be included as an optimization objective and minimized. Both approaches have been studied in the literature and require corresponding modifications to the FBSM algorithm (see [9, 21] for details). Here, we propose a combination of these ideas as a potential direction for future research. Specifically, we may prescribe a target tumour burden while leaving the terminal time free, and seek a treatment strategy that minimizes the cost functional while also minimizing the treatment duration. In other words, our problem takes the following form: for a given *x*(*t*_0_) = (*T*_0_, *C*_0_, *B*_0_, *A*_0_)^T^ and *T* (*τ* ) = *ε* (for a small enough *ε*), minimize

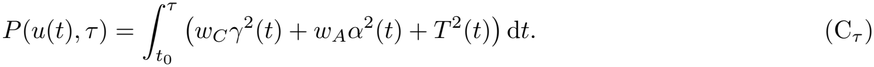

Solving this problem would require the development of a modified FBSM algorithm. Its derivation, theoretical analysis, and convergence properties are beyond the scope of the present work. Nevertheless, we believe that investigating this formulation may provide a valuable direction for future research.

There have been some recent works using optimal control for cancer therapies, such as [13]. However, there the authors use a linear running cost hence the obtained optimal control is bang-bang. The paper [12] uses both linear and quadratic cost functions to assess optimal control for leukemia, however there is only one control function for the chemotherapy. None of those, or other papers in the literature have considered the presence of saboteur bacteria and consequently the combined antibiotic-chemotherapeutic treatment.

Additional future research directions include extending the model with pharmacokinetics and pharmaco-dynamics, and more involved cancer growth submodels. Validation using real experimental data would also be desirable, however biological data for cancer-bacteria systems with therapies are still very scarce, and limited to some very special situations ([6]).

## CRediT authorship contribution statement

Anna Geretovszky: Conceptualization, Formal analysis, Methodology, Visualization, Writing – original draft, Writing – review & editing.

Gergely Röst: Conceptualization, Formal analysis, Funding acquisition, Methodology, Supervision, Writing – review & editing.

## Declaration of Competing Interest

The authors declare that they have no known competing financial interests or personal relationships that could have appeared to influence the work reported in this paper.

## Acknowledgement

This study was supported by the National Research, Development and Innovation Fund, Hungary, grant number 2022–2.1.1-NL-2022–00005.

## Notes

### Competing Interest Statement

The authors have declared no competing interest.

